# Heterologous expression and cell membrane localization of dinoflagellate rhodopsins in mammalian cells

**DOI:** 10.1101/2020.05.03.075168

**Authors:** Minglei Ma, Xinguo Shi, Senjie Lin

## Abstract

Rhodopsins are now found in all domains of life, and are classified into two large groups: type II, found in animals and type I found in microbes including Bacteria, Archaea, and Eukarya. While type II rhodopsin functions in many photodependent signaling processes including vision, type I among others contains rhodopsins that function as a light-driven proton pump to convert light into ATP as in proteobacteria (named proteorhodopsin). Proteorhodopsin homologs have been documented in dinoflagellates, but their subcellular localizations and functions are still poorly understood. Even though sequence analyses suggest that it is a membrane protein, experimental evidence that dinoflagellate rhodopsins are localized on the plasma membrane or endomembranes is still lacking. As no robust dinoflagellate gene transformation tool is available, we used HEK 293T cells to construct a mammalian expression system for two dinoflagellate rhodopsin genes. The success of expressing these genes in the system shows that this mammalian cell type is suitable for expressing dinoflagellate genes. Immunofluorescence of the expressed protein locates these dinoflagellate rhodopsins on the cell membrane. This result indicates that the protein codons and membrane targeting signal of the dinoflagellate genes are compatible with the mammalian cells, and the proteins’ subcellular localization is consistent with proton pump rhodopsins.

## Introduction

The energy that supports life activities in the ocean ultimately comes from sunlight, via photosynthesis as the main energy conversion mechanism. Rhodopsin is a newly discovered protein in microbial organisms, the proton-pump type of which can produce proton gradients thus facilitate ATP generation. (Béja et al. 2000; Grote et al. 2014), the main energy currency of cells, without the involvement of photosynthetic apparatus. It has been estimated that 15%-70% bacteria living in surface ocean carry proton-pump rhodopsin (PR), which are called proteorhodopsins, from the tropical Red sea to the Arctic at different depths (Campbell et al. 2008; Sabehi et al. 2005). In the PR-harboring microbial organisms, the photoreactive chromophore (all-trans retinal) in the PR molecules absorbs light and undergoes conformational changes, resulting in proton generation and efflux across the membrane, the potential to generate ATP upon the action of ATP synthases (Govorunova et al. 2017; Grote et al. 2014). Based on amino acid sequences, rhodopsins similar to that encoding bacterial PR have been found in diverse species of dinoflagellates (Lin et al. 2010). These dinoflagellate homologs share conserved residues with bacteria PR, including retinal pocket, proton donor and receptor, and light tuning (Lin et al. 2010; Shi et al. 2015). Shi et al. (2015) found that PR in the dinoflagellate, *Prorocentrum donghaisense*, exhibited a strong diel rhythm in transcript abundance: high in the light period and low in the dark. In another dinoflagellate, *Oxyrrhis marina*, PR genes were found to be up-regulated while being starved and illuminated relative to being starved and placed under darkness, suggestive of a role as an energy supplementing mechanism to enhance starvation survival (Guo et al. 2014).

However, the function of dinoflagellate PR homologs remains unknown and even the subcellular localization of dinoflagellate rhodopsin is not so clear. Slamovits et al. (2011) used specific antibody to study subcellular localization of PR in *O. marina* and found that it was localized in cytoplasmic endomembrane rather than plasma membrane as in bacteria. Yet most of the PR homologs in the diverse dinoflagellates have not been investigated in this regard. Homologous transformation of the gene (i.e. in native species) would be most straightforward approach to determining their localizations; unfortunately, a robust transformation tool is not yet available. Taking advantage of well-developed transformation technology in animal cells, we transfected PR genes of two dinoflagellates, *P. donghaisense* (Pd-PR) and *Alexandrium catenella* (Ac-PR), into HEK 293T cells using the Piggybac system (Wilson et al. 2007). After transfection and selection under antibiotic pressure, the stable transformed cells were harvested, total RNA extracted, and reverse transcription-PCR performed to detect expression of the inserted PR genes. Furthermore, Western blot was conducted to verify successful translation in the 293T cells. Finally, subcellular localization was analyzed, which showed that the expressed dinoflagellate PRs are targeted to the plasma membrane of the 293T cells.

## Results

### Detecting expression of the inserted gene cassette via GFP fluorescence

Because the expression vector contained a GFP gene, successful transformation and expression of the gene cassette can be easily observed under an epifluorescence microscope. After selection under puromycin, we found that every cell in the transformed cell cultures expressed GFP (Fig. 2), indicating successful transfection and expression of the gene cassette.

**Fig. 1.**
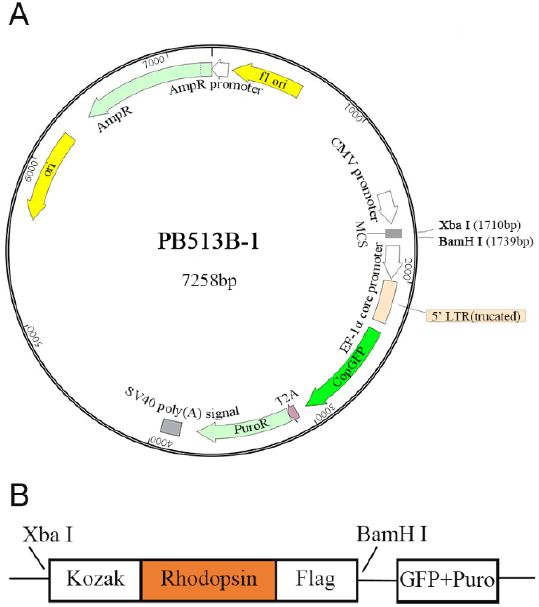
PB513b-1 vector (**a**) and PB513b-Pd/Ac-PR vector (**b**) used in this study. Kozak sequence (GCCACC was added to promote expression. Flag is a tag (GATTACAAGGATGACGACGATAAG) which encodes a short peptide, against which antibodies are available for detection. GFP, green fluorescence protein. Puro, a puromycin (antibiotic) resistance gene to facilitate selection of transformed cells from untransformed cells (lacking the resistance gene)

**Fig. 2.**
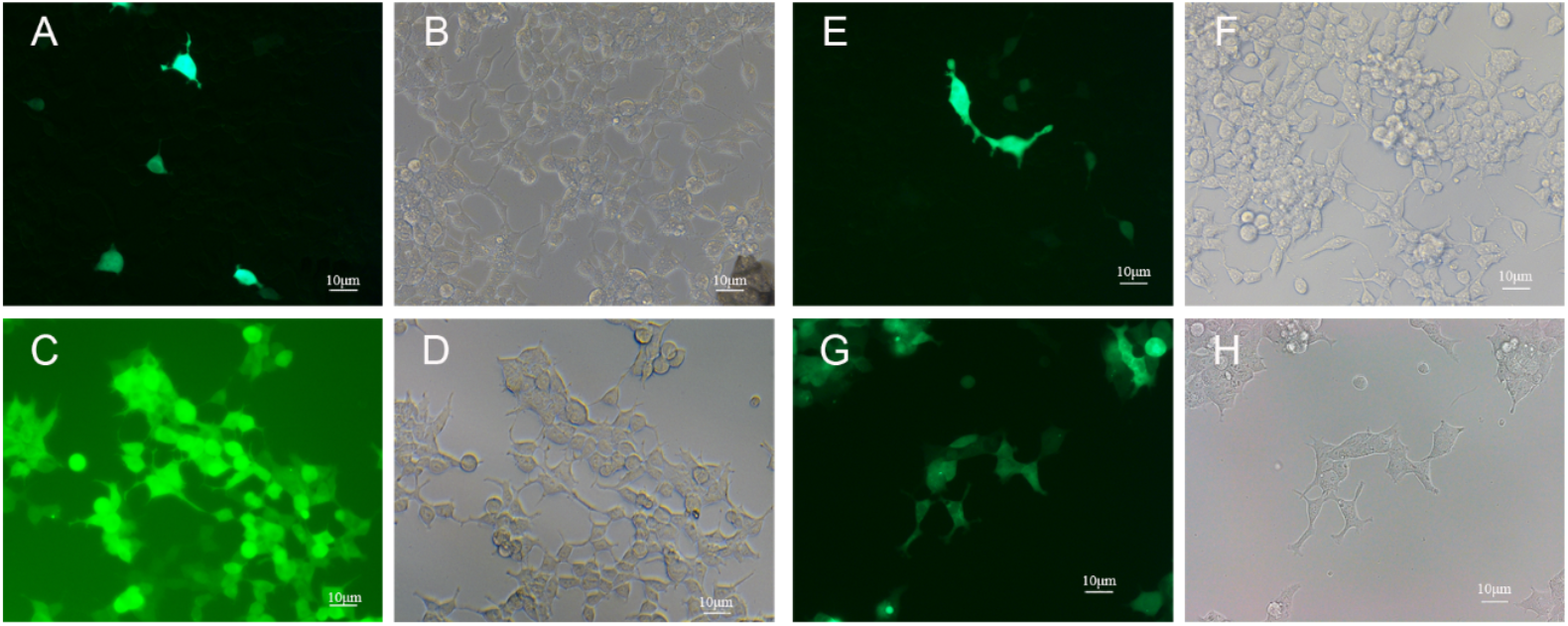
Results of selection by puromycin. **a, b** Microscopic images of 239T cells after being co-transfected with vectors, PB513b-Pd-PR and PB200-PA. **c, d** Microscopic images of PB513b-Pd-PR transfected 293T cells after being selected by puromycin. **e, f** Microscopic images after being co-transfected with vectors PB513b-Ac-PR and PB200-PA. **g, h** Microscopic images of PB513b-Ac-PR transfected 293T cells after being selected by puromycin. Images **a, c, e** and **g** are fluorescence images, λ =470nm (20×). Images **b, f, d** and **h** are bright field (20×)

### Western blot and immunofluorescence to detect expression and localization of dinoflagellate PRs

To determine the expression of Pd-PR and Ac-PR, we performed Western blotting using anti-Flag antibody which was supposed to specifically recognize the Flag tag that was fused with the PR gene at the 3’-end. SDS-PAGE gel electrophoresis was run for total proteins of the Pd-PR and Ac-PR transformed mammalian cells, respectively. On the Western blot, the antibody recognized a single, highly abundant band of the predicted size, 30 kDa (Fig. 3A and Fig. 3B). Because Flag gene was placed downstream of rhodopsin genes, this result indicated the correct expression of both dinoflagellate PR fusion proteins in the mammalian cell expression system. The band detected by anti-Flag antibody reflected the molecular mass of Pd-PR and Ac-PR.

**Fig. 3.**
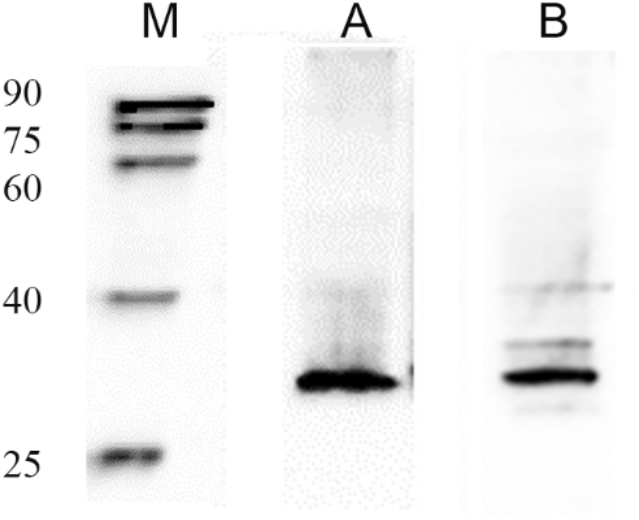
Results of Western blot. **a** Western blot of total protein from 293T cells which was transfected with PB513b-Pd-PR. **b** Western blot of total protein from 293T cells which was transfected with PB513b-Ac-PR. Expected protein size is 30 kDa. **m** Protein Marker. Twenty μg total protein was loaded per lane except for lane m.

In the immunofluorescence analysis, DAPI-staining showed the position of the nucleus in the center of the cells (Fig. 4A). GFP is a cytoplasmic protein and the green fluorescence was localized throughout the 293T cells, as expected (Fig. 4B). In contrast, immunofluorescence of the dinoflagellate PR-Flag fusion protein was localized on the plasma membrane of the cells while the control group exhibited no Flag fluorescence (Fig. 4).

**Fig. 4.**
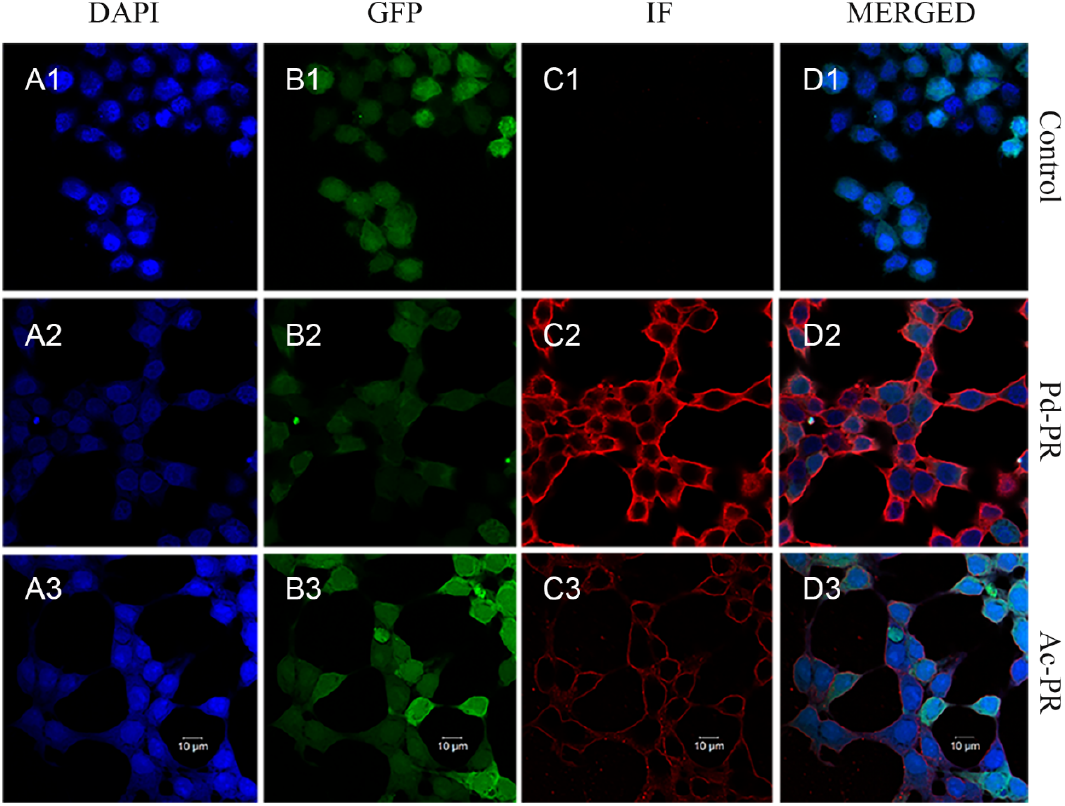
Cell membrane localization of Pd-PR and Ac-PR in 293T cells. The fluorescence microscopic images on the right show localization of Pd-PR and Ac-PR in 293T cells using immunofluorescence assay with the anti-Flag antibody. From **a** to **d**, images show blue fluorescence of DAPI (4,6-diamidino-2-phenylindole) staining of DNA (**a**), green fluorescence of GFP (**b**), red fluorescence of Flag (**c**), and merger for **a, b**, and **c** (**d**). Top row, control group which was transfected with blank PB513b-1 vector. Middle row, 293T cells transfected with PB513b-Pd-PR. Bottom row, 293T cells transfected with PB513b-Ac-PR

## Discussion

To our knowledge, this is the first successful expression of a dinoflagellate gene in a mammalian cell expression system. This system has the advantage that there are well developed protocols and expression vectors. To date, successful nuclear gene transformation for dinoflagellates is still limited (Sprecher et al. 2020) and has yet to emerge for the strongly thecate species such as *P. donghaiense* and *A. catenella*. At this juncture, the development of an effective protocol to expression dinoflagellate genes in the mammalian expression system is valuable.

Microbial rhodopsin heterologous expression has mostly been conducted using *Escherichia coli* cell system (Friedrich et al. 2002; Inoue et al. 2013; Pushkarev et al. 2016; Pushkarev et al. 2018). *E. coli* is a good choice for expressing bacterial PRs. Some researchers have used mammalian neurons to express microbial rhodopsin for optogenetics (i.e. Archaerhodopsin-3, from *Halorubrum sodomense*) to facilitate functional studies via electrophysiological measurements (McIsaac et al. 2015). Interestingly, Kralj et al. (2012) expressed Archaerhodopsin-3(Arch) as a voltage indicator in cultured rat hippo-campal neurons and observed targeting of Arch to the plasma membrane. McIsaac et al. (2014) used 293 cells to express microbial rhodopsins. Our successful expression of two dinoflagellate PRs extend the use of the cell system for microeukaryotic rhodopsin expression, and demonstrate that using mammalian cells to express dinoflagellate proton-pump rhodopsins is feasible. Beside the predicted band, there are several additional faint bands above 30kDa in the results of Western blot (Fig. 3), which most likely represent oligomers of rhodopsin (Ran et al. 2013).

There is no experimental documentation of subcellular localization of PRs in photosynthetic dinoflagellates. The heterotrophic *O. marina* is the only dinoflagellate whose rhodopsin subcellular localization has been examined (Slamovits et al. 2011). Using a PR antibody to detect native PR protein, that study showed that the PR in this species was localized in the cytoplasmic compartment, likely in microsomal membranes, possibly in food vacuole membranes. The results of immunofluorescence analysis in our study showed that the PRs from the two photosynthetic dinoflagellates, *P. donghaiense* and *A. carterae*, when expressed in the HEK 293T cells, were localized in the plasma membrane. The red fluorescence level of Pd-PR is higher than that of Ac-PR (Fig. 4C2, C3), but the bands in result of Western blot (Fig.3) have comparable intensity. It may be because we did the Western blot and Immunofluorescence analysis using different cell samples separately. Therefore, the quantitative difference between Western blot and Immunofluorescence might reflect sample to sample difference or difference in antibody affinity to denatured and native Ac-PR. Based on the subcellular localization computational models CELLO (Yu et al. 2006) and WOLF PSORT (Horton et al. 2007), we used the deduced amino acid sequences of the two dinoflagellate PRs to conduct a prediction. Both models gave highest likelihood of these PRs as plasma membrane proteins. Interestingly, the animal submodel in CELLO gave much stronger plasma membrane localization prediction than did plant and fungus submodels. We further attempted to identify a signal peptide by using SignalP4.1; however, no significant signal peptide was found, as in the case of *O. marina* PR reported by Slamovits et al. (2011). These results suggest that dinoflagellates use some unknown signaling mechanism that is compatible with mammalian protein targeting machinery.

PR in dinoflagellates is highly similar to proteorhodopsin, which are globally abundant in bacteria (Béja et al. 2013; Gómez-Consarnau et al. 2010) and well characterized as a proton pump (Béja et al. 2000; Inoue et al. 2016). Since its first discovery in dinoflagellates (Lin et al. 2010), eukaryotic homologs of PR have also been found in other eukaryotes microorganisms, such as diatoms, haptophytes and cryptophytes (Marchetti et al. 2015), and recently found in arctic microbial eukaryotes (Vader et al. 2018). The dinoflagellate homologs share the highly conserved residues with bacterial PR, including residues to bind retinal, absorbing light spectrum tuning, and proton donor and receptor residues (Lin et al. 2010); this is true for the two homologs studied here, Pd-PR (Shi et al. 2015), and Ac-PR (Supplementary Figure S1). Both proteorhodopsin in bacteria and eukaryotes can enhance organism survival during starvation or nutrient limitation when incubated in the light (Gómez-Consarnau et al. 2010; Guo et al. 2014; Marchetti et al. 2015).

Until now, two methods have been used to investigate the function of rhodopsins as a proton pump: one is based on the color of cloned cells cultured on retinal plates (Martinez et al. 2007), and the other is through the measurement of pH change of cloned cells when exposed to light (Pushkarev et al. 2016). In *O. marina*, the high expression of rhodopsins renders the cells to appear pink (not shown). Heterologous expressing *P. donghaiense* PR in *E. coli*, Shi et al. (2015) showed a rather wide blue to green waveband of absorbance spectrum. Unfortunately, the presence of chloroplasts in the photosynthetic dinoflagellates precludes the possibility to visualize or spectrophotometrically measure the light absorbance. Our attempt to measure pH change using the method previously used to test pH change of proteorhodopsin expressing-*E. coli* under light did not show pH change in the cell suspension of the transfected 293T cell line in response to illumination (results not shown). There are at least two possibilities, one being that the expression level was not adequate to produce measurable change in pH upon illumination, and the second being that these dinoflagellate PRs function other than as proton pumps. These and other possibilities require further experimental studies to examine. Nevertheless, the expression system developed here provides a valuable avenue for easily heterologous expressing dinoflagellate genes and characterizing features such as signal peptide and subcellular localization.

## Materials and Methods

### Construction of mammalian cells expression system

The Piggybac system used in the study included two vectors, PB513b-1(Fig. 1A) and PB200-PA (Miaoling Bio,). In principle, a rhodopsin gene would be inserted into PB513b-1, and PB200-PA would induce transfected mammalian cells to produce transposase, which would “cut” sequences between “TTAA” sites on vector PB513b-1, and “paste” it into “TTAA” sites in genomic DNA. Expression vectors, PB513b-Pd-PR and PB513b-Ac-PR were constructed as follows.

*Prorocentrum donghaisense* and *Alexandrium catenella* cultures were provided by the Center for Collections of Marine Algae in Xiamen University (source culture number: CCMA-364 and CCMA-174, separately). They were grown in L1 medium at 20°C under a 14:10 light dark cycle with a photon flux of 100 micromole m^-2^ s^-1^. Samples were collected in the exponential growth phase and RNA was extracted following the protocol of Trizol protocol coupled with Direct-zol RNA mimiprep columns (Zymol), as previously reported (Shi et al. 2015). The full-length cDNA of Pd-PR genes were amplified by PCR using primers Pd-PR-F (5’-CGCTCTAGAGCCACCATGGTGATGTACCCGATGAGCG-3’) and Pd-PR-R (5’-GGCGGATCCTCACTTATCGTCGTCATCCTTGTAATCAGCAAGCAGGGCCCCATC-3’). Similarly, the full length cDNA of Ac-PR genes were amplified by PCR using primers Ac-PR-F (5’-CGCTCTAGAGCCACCATGGCTCCAATCCCTGATGGT-3’) and Ac-PR-R (5’-GGCGGATCCTCACTTATCGTCGTCATCCTTGTAATCCGACGACATCAGACTGCC-3’). The forward primers contained the optimal Kozak’s sequence (GCCACC) to promote expression (Kozak et al. 1987) and the reverse primers contained the Flag-tag (GATTACAAGGATGACGACGATAAG) (Einhauer et al. 2001) to facilitate detection of expression of the inserted gene. The PR fragment were inserted into the XbaI and BamHI sites of the PB513b-1 transformation vector (Fig. 1B). To facilitate determination of successful transformation and expression of the inserted gene cassette, green fluorescence protein (GFP) gene was placed in the vector, located downstream of the target gene insertion site, but not fused with the target gene to avoid interference with the expression of the target gene. After construction, the inserted sequences in the expression vectors were verified using Sanger sequencing (supplementary). The vectors were used in cell transfection after sequence alignment with Pd-PR sequence (Genbank accession no. KM282617.1) and Ac-PR sequence (Genbank accession no. KF651056.1) and verification of sequence accuracy.

### Cell transfections and selection

Human embryonic kidney (HEK) transformed 293 (293T) cells were obtained from School of Life Science, Xiamen University. 293T cells were grown in Dulbecco modified Eagle medium (DMEM, Hyclone) with 10% fetal bovine serum (FBS, BOVOGEN) and 1% antibiotics (penicillin-streptomycin, Invitrogen) (DMEM-FBS-Pen-Strep) at 37°C in a humidified atmosphere containing 5% CO2.

About 1.5 × 10^5^ cells were transferred into each well of a 24-well-plate one day prior to transfection. For each well, 0.5 μg PB513b-Pd/Ac-PR and 0.5 μg PB200-PA were transfected using Suohua-sofast according to standard protocols (Sunmabio). Empty expression PB513b vector was used as the control group (PB513b-1). One day after transfection, the DMEM-FBS-Pen-Strep was replaced by the medium containing 3 μg/ml puromycin (DMEM-FBS-Pen-Strep-Puro) for 3 days. Then the cells in each well were trypsinized and seeded onto one 10 cm plate in the medium containing 3 μg/ml puromycin for selection. After two weeks, the cells in each plate were trypsinized and inoculated onto a 96 wells plate. After 4 hours, the plates were checked under the microscope to identify wells that contained only one cell, and the single-cell wells were marked. Following 3 days of incubation, the clonal cell population from each of the single-cell wells was transferred to a new 24-well-plate. Then the cells in each well were trypsinized and seeded onto one 10 cm plate in the medium containing 3 μg/ml puromycin to promote growth of the transformed cells.

### Reverse transcription (RT)-PCR

RT-PCR was performed to examine whether the transfected 293T cells successfully expressed the two dinoflagellate PRs. First, total RNA was extracted from the two transformed lines of 293T cells using RNA Miniprep kit (Zymo Research). For cDNA to be used in RT-PCR, total RNA was reverse-transcribed using GoScript Reverse Transcriptase Kit (Promega). Second, PCR was carried out with the primers Pd-PR-F and Pd-PR-R for *P. donghaiense* PR and Ac-PR-F and Ac-PR-R for *A. carterae* PR. PCR was run under the program consisting of initial denaturation at 95°C for 3 min, followed by 30 cycles 95°C 30s, 55°C 45s, 72°C 1min, and a final step of 72°C for 3 min.

### Western blot

After two weeks’ selection by puromycin, cultured cells of the two transformed lines were collected using centrifugation for 5 min at 6280 rad/min and 4°C. The pellet was washed in 1 x PBS and total proteins were extracted in ice-cold RIPA Lysis buffer (Beyotime) for 10 min. Total protein concentrations were determined using BCA assay. The crude extracts were then incubated with 6× SDS at 95°C for 10 min to denature proteins. After centrifugation at 14000*g* for 10 min, the supernatants were loaded onto a 10% SDS-PAGE gel (~20 μg per lane). A protein molecular mass marker was loaded on a separate lane. After electrophoresis at 90V for 30 min, the separated proteins were transferred onto PVDF membranes for 30 min using BioRad Trans-Blot Turbo system (Bio-Rad) at 110V. Pd-PR and Ac-PR were detected using antiFlag antibody with clarity western ECL substrate kit (Bio-Rad).

### Immunofluorescence analysis to determine subcellular localization

Puromycin selected cells were used for microscopic immunofluorescence analysis. Cells grown on coverslips coated with polyethyleneimine (NEST) at multi-well plates. Coverslips were collected and washed in 1 x PBS for three times then fixed with 4% formaldehyde for 30 min. The cells were rinsed three times with 1 x PBS, blocked with 5% BSA in PBS for 30 min, and then incubated overnight at 4°C with the anti-Flag antibody diluted to 1:50 with PBS. After three rinses with PBS, the cells were incubated in goat anti mouse IgG(H+L) (EarthOx) diluted 1:200 with PBS for 1h at room temperature, and rinsed with PBS for 3 times. Then cells were incubated with DAPI in darkness for 5 min to counter stain the nucleus and rinsed with PBS for 4 times. Immunofluorescence of the cells was viewed using multiphoto laser scanning microscopy (LSM780 NLO) and images were acquired with the Zen software (Zeiss, Oberkochen, Germany).

## Acknowledgements

We wish to thank professor Kejian Wang for generously allowing us to use his cell culture platform. We also thank all members of Marine EcoGenomics Laboratory of Xiamen University, China for various ways of assistance in this study. The work was supported by the National Key Research and Development Program grant 2017YFC1404302 and Natural Science Foundation of China grants NSFC 31661143029 and 41776116.

## Authors’ Contribution

MLM performed the experiments, analyzed the data, wrote the paper, and prepared the figures; XGS provided sample, providing advice on the experiment, and revised the manuscript; SJL conceived and supervised the design of the project and revised the manuscript.

## Compliance with Ethical Standards

### Conflicts of Interest

we declare that we have no conflicts of interest.

### Animal and human rights statement

This article does not contain any studies with human participants or animals performed by any of the authors

## Notes

### Competing Interest Statement

The authors have declared no competing interest.

### Summary of Updates

Correction of typos of Type I and Type II in the Abstract.

## Reference

Beja O, Aravind L, Koonin EV, Suzuki MT, Hadd A, Nguyen LP, Jovanovich SB, Gates CM, Feldman RA, Spudich JL, Spudich EN, DeLong EF (2000). Bacterial rhodopsin: evidence for a new type of phototrophy in the sea. Science 289:1902–1906

Béjà O, Pinhassi J, Spudich JL (2013). Proteorhodopsins: widespread microbial light-driven proton pumps. In Levin SA (ed), Encyclopedia of biodiversity, 2nd ed, pp 280–285.

Campbell BJ, Waidner LA, Cottrell MT, Kirchman DL (2008). Abundant proteorhodopsin genes in the North Atlantic Ocean. Environ Microbiol 10:99–109

Einhauer A, Jungbauer A (2001). The FLAG™ peptide, a versatile fusion tag for the purification of recombinant proteins. J Biochem Biophys Meth 49:455–465

Friedrich T, Geibel S, Kalmbach R, Chizhov I, Ataka K, Heberle J, Engelhard M, Bamberg E (2002) Proteorhodopsin is a light-driven proton pump with variable vectoriality. J Mol Biol 321:821–838

Gómez-Consarnau L, Akram N, Lindell K, Pedersen A, Neutze R, Milton DL, González JM, Pinhassi J (2010) Proteorhodopsin phototrophy promotes survival of marine bacteria during starvation. PLoS Biol 8:e1000358

Govorunova EG, Sineshchekov OA, Li H, Spudich JL (2017) Microbial rhodopsins: diversity, mechanisms, and optogenetic applications. Ann Rev Biochem 86:845–872

Grote M, Engelhard M, Hegemann P (2014) Of ion pumps, sensors and channels—perspectives on microbial rhodopsins between science and history. Biochim Biophys Acta 1837:533–545

Guo Z, Zhang H, Lin S (2014). Light-promoted rhodopsin expression and starvation survival in the marine dinoflagellate *Oxyrrhis marina*. PLoS ONE 9:e114941

Horton P, Park KJ, Obayashi T, Fujita N, Harada H, Adams-Collier CJ, Nakai K (2007) WoLF PSORT: protein localization predictor. Nucleic Acids Res 35:W585–W587

Inoue K, Ono H, Abe-Yoshizumi R, Yoshizawa S, Ito H, Kogure K, Kandori H (2013) A light-driven sodium ion pump in marine bacteria. Nat Commun 4:1678

Inoue K, Ito S, Kato Y, Nomura. Y, Shibata M, Uchihashi T, Tsunoda SP, Kandori H (2016) A natural light-driven inward proton pump. Nat Commun 7:13415

Kozak M (1987) At least six nucleotides preceding the AUG initiator codon enhance translation in mammalian cells. J Mol Biol 196:947–950

Kralj JM, Douglass AD, Hochbaum DR, Maclaurin D, Cohen AE (2012) Optical recording of action potentials in mammalian neurons using a microbial rhodopsin. Nat Methods 9:90

Lin S, Zhang H, Zhuang Y, Tran B, Gill J (2010) Spliced leader-based metatranscriptomic analyses lead to recognition of hidden genomic features in dinoflagellates. Proc Natl Acad Sci USA 107:20033–20038

Martinez A, Bradley AS, Waldbauer JR, Summons RE, DeLong EF (2007) Proteorhodopsin photosystem gene expression enables photophosphorylation in a heterologous host. Proc Natl Acad Sci USA 104:5590–5595

Marchetti A, Catlett D, Hopkinson BM, Ellis K, Cassar N (2015) Marine diatom proteorhodopsins and their potential role in coping with low iron availability. ISME J 9:2745–2748

McIsaac RS, Bedbrook CN, Arnold FH (2015) Recent advances in engineering microbial rhodopsins for optogenetics. Curr Opin Struct Biol 33:8–15

McIsaac RS, Engqvist MK, Wannier T, Rosenthal AZ, Herwig L, Flytzanis NC, Imasheva ES, Lanyi JK, Balashov SP, Gradinaru V, Arnold FH (2014) Directed evolution of a far-red fluorescent rhodopsin. Proc Natl Acad Sci USA 111:13034–13039

Pushkarev A, Béjà O (2016) Functional metagenomic screen reveals new and diverse microbial rhodopsins. ISME J 10:2331

Pushkarev A, Hevroni G, Roitman S, Shim JG, Choi A, Jung KH, Béjà O (2018) The use of a chimeric rhodopsin vector for the detection of new proteorhodopsins based on color. Front Microbiol 9:439

Ran T, Ozorowski G, Gao Y, Sineshchekov OA, Wang W, Spudich JL, Luecke H (2013). Cross-protomer interaction with the photoactive site in oligomeric proteorhodopsin complexes. Acta Crystallogr 69:1965–1980

Sabehi G, Loy A, Jung KH, Partha R, Spudich JL, Isaacson T, Hirschberg J, Béjà O (2005) New insights into metabolic properties of marine bacteria encoding proteorhodopsins. PLoS Biol 3: e273

Shi X, Li L, Guo C, Lin X, Li M, Lin S. (2015) Rhodopsin gene expression regulated by the light dark cycle, light spectrum and light intensity in the dinoflagellate Prorocentrum. Front Microbiol 6:555

Slamovits CH, Okamoto N, Burri L, James ER, Keeling PJ (2011) A bacterial proteorhodopsin proton pump in marine eukaryotes. Nat Commun 2:183

Sprecher BN, Zhang H, Lin S (2020) Nuclear Gene Transformation in the Dinoflagellate *Oxyrrhis marina*. Microorganisms 8:126

Vader A, Laughinghouse IV HD, Griffiths C, Jakobsen KS, Gabrielsen TM (2018) Proton-pumping rhodopsins are abundantly expressed by microbial eukaryotes in a high-Arctic fjord. Environ Microbiol 20:890–902

Wilson MH, Coates CJ, George Jr AL (2007) PiggyBac transposon-mediated gene transfer in human cells. Mol Ther 15:139–145

Yu C-S, Chen Y-C, Lu C-H, Hwang J-K (2006) Prediction of protein subcellular localization. Proteins 64:643–651

